# Prioritizing putative influential genes in early life cardiovascular disease susceptibility by applying tissue-specific Mendelian randomization

**DOI:** 10.1101/298687

**Authors:** Kurt Taylor, George Davey Smith, Caroline L Relton, Tom R Gaunt, Tom G Richardson

**Author notes:** Corresponding author: Dr. Tom G. Richardson, MRC Integrative Epidemiology Unit, Bristol Medical School (Population Health Sciences), University of Bristol, Oakfield House, Oakfield Grove, Bristol BS8 2BN, UK.

## Abstract

**Background:** The extent to which changes in gene expression can influence cardiovascular disease risk across different tissue types has not yet been systematically explored. We have developed an analytical framework that integrates tissue-specific gene expression, Mendelian randomization and multiple-trait colocalization to develop functional mechanistic insight into the causal pathway from genetic variant to complex trait.

**Methods:** We undertook a transcriptome-wide association study in a population of young individuals to uncover genetic variants associated with both nearby gene expression and cardiovascular traits. Two-sample Mendelian randomization was then applied using large-scale datasets to investigate whether changes in gene expression within certain tissue types may influence cardiovascular trait variation. We subsequently performed Bayesian multiple-trait colocalization to further interrogate findings and also gain insight into whether DNA methylation, as well as gene expression, may play a role in disease susceptibility.

**Results:** Eight genetic loci were associated with changes in gene expression and early life measures of cardiovascular function. Our Mendelian randomization analysis provided evidence of tissue-specific effects at multiple loci, of which the effects at the *ADCY3* and *FADS1* loci for body mass index and cholesterol respectively were particularly insightful. Multiple trait colocalization uncovered evidence which suggested that changes in DNA methylation at the promoter region upstream of *FADS1/TMEM258* may also play a role in cardiovascular trait variation along with gene expression. Furthermore, colocalization analyses were able to uncover evidence of tissue-specificity, most prominantly between *SORT1* expression in liver tissue and cholesterol levels.

**Conclusions:** Disease susceptibility can be influenced by differential changes in tissue-specific gene expression and DNA methylation. Our analytical framework should prove valuable in elucidating mechanisms in disease, as well as helping prioritize putative causal genes at associated loci where multiple nearby genes may be co-regulated. Future studies which continue to uncover quantitative trait loci for molecular traits across various tissue and cell types will further improve our capability to understand and prevent disease.

## Introduction

Despite recent efforts in research and development, cardiovascular disease still poses one of the greatest threats to public health throughout the world, accounting for more deaths than any other cause [1]. Since their development, genome-wide association studies (GWAS) have identified thousands of different genetic loci associated with complex disease traits [2]. An example of their successful application within cardiovascular research is the identification of numerous genetic variants associated with low density lipoprotein (LDL) cholesterol levels [3], which is a causal mediator along the coronary heart disease progression pathway [4,5]. However, the functional and clinical relevance for the vast majority of GWAS results are still unknown, emphasizing the importance of developing our understanding of the causal pathway from single nucleotide polymorphism (SNP) to disease.

A large proportion of associations detected by GWAS are located in non-coding regions of the genome [6], suggesting that the underlying SNPs influence complex traits via changes in gene regulation [7]. Recent efforts have incorporated messenger ribonucleic acid (mRNA) expression data into analyses to determine whether SNPs identified by GWAS influence levels of gene expression (i.e. whether they are expression quantitative trait loci [eQTL]) as well as complex traits [8]. Novel methods have integrated eQTL data with summary association statistics from GWAS [9] to identify genes whose nearby (*cis*) regulated expression is associated with traits of interest (widely defined as variants within 1 megabase (Mb) on either side of a genes transcription start site [TSS]) [10]. These types of studies have been referred to as transcriptome-wide association studies (TWAS).

A recent paper has highlighted some limitations that may be encountered by studies integrating transcriptome data to infer causality [11], such as intra-tissue variability and co-regulation amongst proximal genes, making it challenging to disentangle putative causal genes for association signals. This exemplifies the importance of developing methods that investigate tissue-specificity and co-regulation of association signals detected by TWAS. Therefore, there needs to be further research into the most appropriate manner to harness eQTL data (across multiple tissue and cell types) in order to improve the biological interpretation of GWAS findings.

We have developed a systematic framework which can be used to evaluate five potential scenarios that can help explain findings from TWAS (Figure 1). Firstly, we identify putative causal genes responsible for observed association signals, by evaluating the association between lead SNPs and proximal gene expression using eQTL data from the Framingham Heart Study (n=5,257) [8]. We then investigate the relationship between gene expression and complex traits at loci of interest by applying the principles of Mendelian randomization (MR); a method which uses genetic variants associated with an exposure as instrumental variables to infer causality among correlated traits [12,13]. A recent development in this paradigm is two-sample MR, by which effect estimates on exposures and outcomes are derived from two independent datasets, allowing researchers to exploit findings from large GWAS consortia [14]. Applying this approach can therefore be used to help infer whether changes in gene expression (our exposure) may influence a complex trait identified by GWAS (our outcome). Furthermore, as tissue-specificity is fundamental in understanding causal mechanisms involving gene expression, we have used data from the genotype tissue expression project (GTEx) [15] in a number of tissues that could be important in cardiovascular disease susceptibility (Additional file 2: Table S1) to try and disentangle co-regulation amongst proximal genes (i.e. differentiating between scenarios 1, 2 and 3). We refer to this approach as tissue-specific MR, which should prove increasingly valuable in investigating both the determinants and consequences of changes in tissue-specific gene expression as sample sizes increase [12].

**Figure 1.**
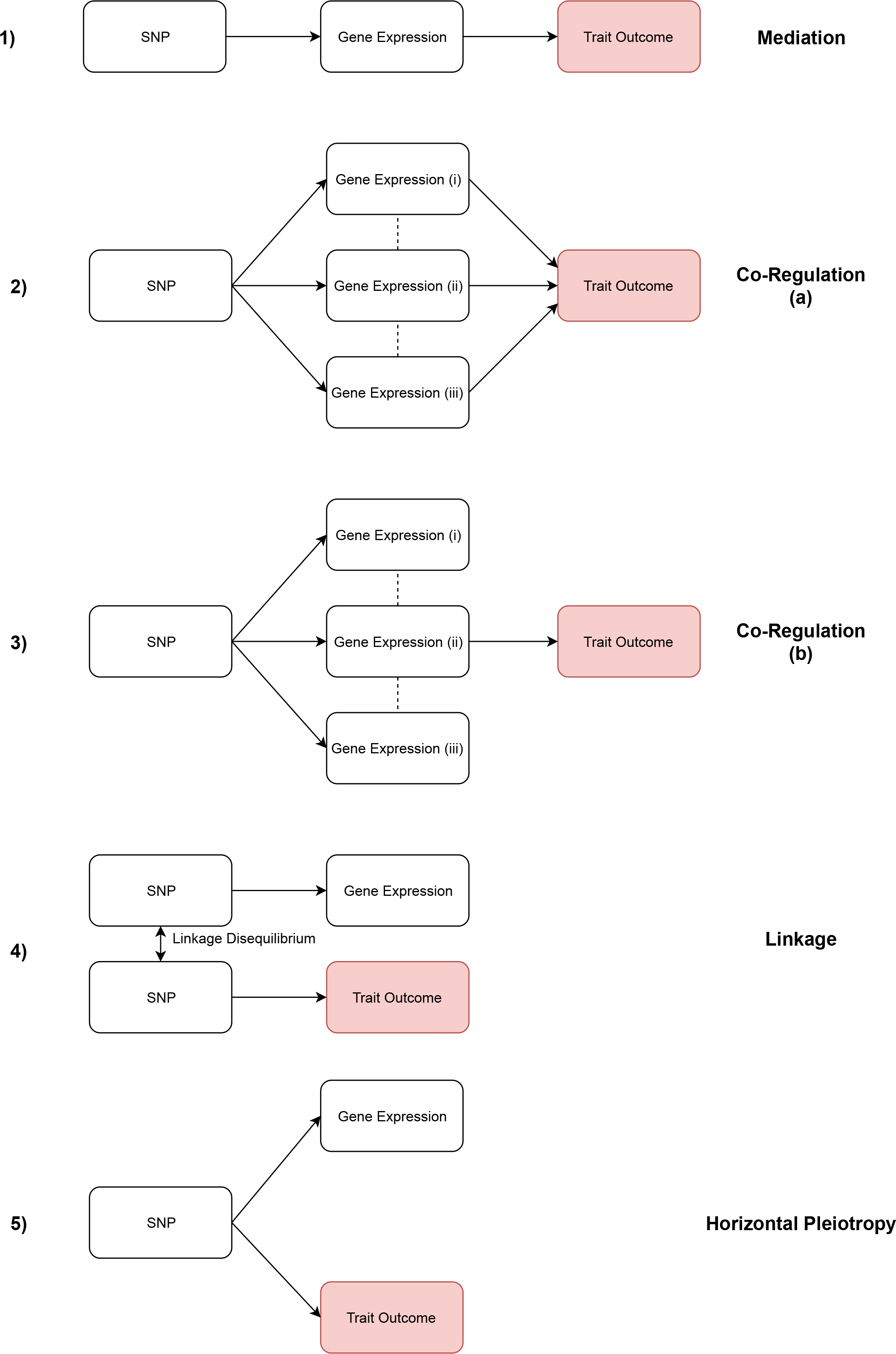
Explanations for observed associations between SNPs and traits. 1) The genetic variant influences the trait, mediated by the expression of a single gene at a locus. 2) The genetic variant influences the trait via multiple genes which are co-regulated with one another. 3) The genetic variant influences the trait via a single gene which is co-regulated with other non-causal genes. 4) The genetic variant that influences the trait is in linkage disequilibrium with another variant which is responsible for changes in gene expression that does not affect the trait. 5) The genetic variant influences both gene expression and the trait outcome by two independent biological pathways (horizontal pleiotropy).

We subsequently apply colocalization analyses [16] at each locus of interest to evaluate whether the same underlying genetic variant is responsible for changes in both gene expression and complex trait, or whether association signals may be a product of linkage disequilibrium (LD) between two causal variants (scenario 4). This analysis can also complement findings from the MR analysis, particularly given that the majority of genes can only be instrumented with a single eQTL using GTEx data. In addition, there has been recent interest in the impact that DNA methylation may have on cardiovascular disease risk via modifications in gene expression [17]. Therefore, we apply multiple-trait colocalization (moloc) [16] at each locus to simultaneously investigate whether the same underlying genetic variant is driving the observed effect on all three traits of interest (i.e. the cardiovascular trait, gene expression and DNA methylation).

Uncovering evidence suggesting that DNA methylation and gene expression may be working in harmony to influence complex traits can improve the reliability of causal inference in this field, as it suggests there may be underlying mechanisms which are consistent with causality (i.e. DNA methylation acting as a transcriptional repressor). However, a major challenge in this paradigm is the lack of accessible tissue-specific DNA methylation/mQTL data akin to GTEx for gene expression. Previous studies have investigated the potential mediatory role of DNA methylation between genetic variant and gene expression using eQTL and mQTL data derived from blood which may act as a proxy for other tissue types [18–20]. Moreover, other studies have demonstrated a surprisingly high rate of replication between mQTL derived from blood and more relevant tissue types for a complex trait of interest [21]. We have therefore undertaken moloc analyses using eQTL derived from both blood and cardiovascular-specific tissue types. Finally, it is also important to note that, along with other approaches which apply causal methods to molecular data, we are currently unable to robustly differentiate mediation from horizontal pleiotropy (scenario 5) [12,22]. However, within this framework we will be able to accommodate additional eQTL as instrumental variables derived from future larger studies in order to address this.

In this study, we demonstrate the value of our framework by applying it to data from the Avon Longitudinal Study of Parents and Children (ALSPAC) using early life measures of cardiovascular function as outcomes. Evaluating putative causal mechanisms apparent early in the life course can be extremely valuable for disease prevention and healthcare, particularly given that cardiovascular disease such as atherosclerosis has been shown to develop in childhood [23]. Therefore, we used ~19,000 cis-eQTL’s observed in adults at risk of cardiac events from the Framingham Heart Study [8] for our TWAS to ascertain whether they influence these cardiovascular traits in young individuals (age ≤ 10). We have further evaluated results using our framework by harnessing summary statistics from large-scale GWAS to demonstrate the value of our approach and validate findings in independent samples.

## Methods

### The Avon Longitudinal Study of Parents and Children (ALSPAC)

Detailed information about the methods and procedures of ALSPAC is available elsewhere [24–26]. In brief, ALSPAC is a prospective birth cohort study which was devised to investigate the environmental and genetic factors of health and development. In total, 14,541 pregnant women with an expected delivery date of April 1991 and December 1992, residing in the former region of Avon, UK were eligible to take part. Participants attended regular clinics where detailed information and bio-samples were obtained. The study website contains details of all the data that is available through a fully searchable data dictionary [27]. All procedures were ethically approved by the ALSPAC ethics and Law Committee and the Local Research Ethics Committees. Written informed consent was obtained from all participants.

#### Genetic data

All children were genotyped using the Illumina HumanHap550 quad genome-wide SNP genotyping platform. Samples were removed if individuals were related or of non-European genetic ancestry. Imputation was performed using Impute V2.2.2 against a reference panel from 1000 genomes [28] phase 1 version 3 [29]. After imputation, we filtered out variants and kept those with an imputation quality score ≥ 0.8 and minor allele frequency (MAF) > 0.01.

#### Phenotypes

The methods and procedures to acquire data for the 14 phenotypes analyzed in this study are as follows. All measurements were obtained at the ALSPAC clinic. Height and weight were measured at age 7 (mean age: 7.5, range: 7.1-8.8). Height was measured to the nearest 0.1 cm with a *Harpenden* stadiometer (*Holtain Crosswell*), and weight was measured to the nearest 0.1 kg on *Tanita* electronic scales. Body mass index (BMI) was calculated as (weight [kg]/(height[m]^2^). Non-fasting blood samples were taken at age 10 (mean age: 9.9, range: 8.9-11.5). The methods on the assays performed on these samples which included total cholesterol, high-density lipoprotein cholesterol, LDL cholesterol (calculated using the Friedewald equation [30]), very low density lipoprotein (VLDL) cholesterol, triglycerides, Apolipoprotein A1 (ApoA1), Apolipoprotein B (ApoB), fasting glucose, fasting insulin, adiponectin, leptin, C-reactive protein (CRP) and interleukin 6 (IL-6) have been described previously [31].

### The Framingham Heart Study

We identified over 19,000 pruned lead cis-eQTLs from Joehanes et al [8] who provide in depth details of the Framingham Heart study and their analysis plan in their paper. Trans-eQTLs were not considered for our analysis to reduce the likelihood of horizontal pleiotropy influencing our findings and also to reduce the burden of multiple testing [32]. This eQTL data was chosen for the initial analysis in ALSPAC due to the larger sample size of transcriptome data from the Framingham Heart Study (n=5,257) using whole blood in comparison to GTEx sample sizes for other tissue types. This allowed us to maximise statistical power to detect association signals which we were then subsequently able to evaluate in detail using data from other tissue types.

### The Genotype-Tissue Expression (GTEx) project

GTEx is a unique open-access online resource with gene expression data for 449 human donors (83.7% European American and 15.1% African American) across 44 tissues. Sample sizes vary between tissues, thus affecting statistical power to identify eQTL. In depth information on the materials and methods for GTEx is available in the latest publication [15]. In short, RNA sequencing samples were sequenced to a median depth of 78 million reads. This is suggested to be a credible depth to quantify accurately genes that may have low expression levels [33]. DNA was genotyped at 2.2 million sites and imputed to 12.5 million sites. We used GTEx eQTL data in all downstream analysis following the discovery analysis in ALSPAC (i.e. Mendelian randomization and multiple-trait colocalization).

### Statistical analysis

Data from ALSPAC were initially cleaned using STATA [version 15] and outliers defined as ± 4 standard deviations from the mean were removed. We plotted histograms to check the data for normality and where necessary applied log-transformation. Using PLINK [version 1.9] [34,35], we undertook an age and sex adjusted TWAS to evaluate the association between *cis*-eQTLs known to influence gene expression and cardiovascular traits. We applied a Bonferroni correction to account for multiple testing which equated to 0.05/the total number of tests undertaken. Using a script derived from the qqman R package [36], results were plotted using a Manhattan plot. We undertook fine mapping across the region 1Mb either side of each lead SNP identified from our TWAS using FINEMAP [37] software. We used the default setting which outputs a maximum of 5 putative causal variants.

#### Tissue-specific Mendelian randomization analysis

To investigate potential causal genes at association signals detected in our TWAS, we applied the principles of MR using the wald method [38] (Additional File 1: Figure S1) to assess whether changes in tissue-specific gene expression (eQTLs as instrumental variables) may be responsible for effects on associated traits. Furthermore, it can help discern whether multiple proximal genes at a region are contributing to trait variation or whether they are likely just co-regulated with causal genes in accessible tissue types such as whole blood, i.e. scenario 3. Firstly, for each lead eQTL from the TWAS we used tissue-specific data from GTEx to discern whether they were cis-eQTL for genes in tissue types which may play a role in the pathology of cardiovascular disease (P < 1 × 10^−4^). If this was not possible then we used eQTL for all genes within a 1MB distance of the lead eQTL. The tissue types evaluated were; adipose - subcutaneous, adipose - visceral (omentum), liver, pancreas, artery - coronary, artery - aorta, heart - atrial appendage and heart - left ventricle. The mean donor age for all tissues included in this analysis resided in the range of 50-55 years. In addition to this, we ran an additional analysis for the association with BMI but investigating effects in the following brain tissues: pituitary, anterior cingulate cortex (BA24) and frontal cortex (BA9).

For this analysis, we used data from large-scale GWAS; A full list of these with details can be found within additional file 2 (Table S2) [39–41]. We then undertook a validation analysis using our ALSPAC data. As cardiovascular trait data is therefore obtained at an earlier stage in the life course compared to the tissue-specific expression data, any associations detected in the validation analysis suggest genetic liability to cardiovascular risk via changes in gene expression. These analyses were undertaken using the MR-Base platform [42]. The only trait we were unable to assess in our analysis was interleukin-6, due to the lack of GWAS summary statistics for this trait. Nonetheless, we still performed MR for the IL-6 data we possessed in ALSPAC. We applied a multiple testing threshold to the MR results to define significance (p<0.05/54). We plotted the results from the validation analysis using volcano plots from the ggplot2 package in R [43]. We also applied the Stieger directionality test [44] to discern whether our exposure (i.e. gene expression) was influencing our outcome (i.e. our complex trait) as opposed to the opposite direction of effect.

#### Multiple-trait colocalization (moloc)

Blood samples were obtained from 1,018 ALSPAC mothers as part of the accessible resource for integrated epigenomics studies (ARIES) [45] from the ‘Focus on Mothers 1’ time point (mean age = 47.5). Epigenome-wide DNA methylation was derived from these samples using the Illumina HumanMethylation450 (450K) BeadChip array. From this data, we obtained effect estimates for all genetic variants within a 1MB distance of lead eQTL from the TWAS and proximal CpG sites (again defined as < 1MB). We then used the moloc [16] method to investigate 2 questions:

1. Is the same underlying genetic variant influencing changes in both proximal gene expression and cardiovascular trait (i.e. investigating scenario 4)
2. Does the genetic variant responsible for these changes also appear to influence proximal DNA methylation levels, suggesting that changes in this molecular trait may also play a role along the causal pathway to disease.

As such, at each locus we applied moloc using genetic effects on 2 different molecular phenotypes (gene expression and DNA methylation (referred to as eQTL and mQTL respectively) along with the associated cardiovascular trait from our GWAS summary statistics. Since we included three traits (i.e. gene expression, DNA methylation and cardiovascular trait), moloc computed 15 possible configurations of how the traits are shared: detailed information on how these are calculated can be found in the original moloc paper [16]. For each independent trait-associated locus, we extracted effect estimates for all variants within 1MB distance of the lead TWAS hit, for all molecular phenotypes and relevant cardiovascular GWAS traits. We subsequently applied moloc in a gene-centric manner, by mapping CpG sites to genes based on the 1 MB regions either side of our TWAS hit. Moloc was subsequently applied to all gene-CpG combinations within each region of interest. We ran this analysis twice, once using expression data from whole blood and again using expression data from a tissue type which was associated with the corresponding trait in the tissue-specific MR analysis (Additional file 2: Table S3).

Only regions with at least 50 SNPs (MAF >= 5%) in common between all three datasets (i.e. gene expression, DNA methylation and cardiovascular trait) were assessed by moloc based on recommendations by the authors. We computed summed PPAs for all scenarios where GWAS trait and gene expression colocalized. When summed PPAs were >= 80%, we reported findings as evidence that genetic variation was influencing cardiovascular traits via changes in gene expression. Furthermore, when summed PPAs relating to DNA methylation were >=80%, there was evidence that DNA methylation may also reside on the causal pathway to complex trait variation via changes in gene expression. In all analyses we used prior probabilities of 1e-04, 1e-06, 1e-07 and 1e-08 as recommended by the developers of moloc based on their simulations [16].

## Results

### Identifying putative causal genes for measures of early life cardiovascular function

We carried out 273,742 tests to valuate the association between previously identified cis-eQTLs [8] with 14 cardiovascular traits in turn within ALSPAC (19,553 cis-eQTLs x 14 traits). After multiple-testing corrections, we identified 11 association signals across 8 unique genetic loci which provided strong evidence of association (p < 1.8 ×10^−7^ [Bonferroni corrected threshold: p<0.05/273,742]). These results can be found in Table 1 and are illustrated in Figure 2. The region near *SORT1* was associated with total cholesterol, LDL cholesterol and ApoB. Additionally, the *LPL* region was associated with both triglycerides and VLDL cholesterol.

**Figure 2.**
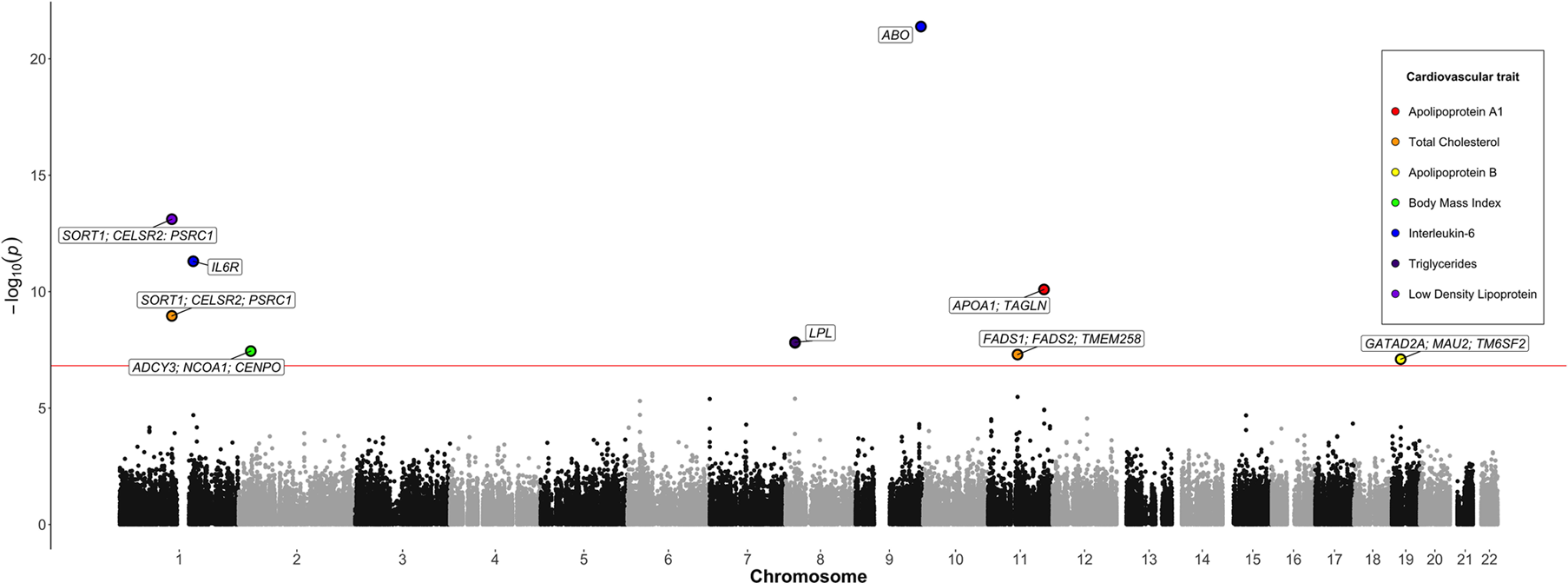
Manhattan plot illustrating observed associations between eQTLs and cardiovascular traits in ALSPAC. Analysed SNPs are plotted on the x-axis ordered by chromosomal position against −log 10 p values which are plotted on the y-axis. SNPs that survived the multiple testing threshold (1.8 × 10^−7^ - represented by the red horizontal line) are coloured according to their associated trait and annotated with potential causal gene symbols.

We undertook fine-mapping 1Mb either side of the lead SNP at each locus identified in our initial analysis to investigate which SNP(s) may be driving the observed effects of complex traits. Posterior probability of association’s (PPA) from FINEMAP [37] suggested that there was most likely only a single variant influencing trait variation for seven of the eleven total loci. For the other four loci, FINEMAP suggested there may be multiple variants influencing traits (Additional file 2: Table S4).

**Table 1.** Results of the TWAS between Genetic Variants Influencing Gene Expression and Cardiovascular Traits in ALSPAC.

|  |  |  |  |  |  |  |
| --- | --- | --- | --- | --- | --- | --- |
| **Tag SNP** | **Gene(s)** | **Trait** | **Sample Size** | **Beta** | **SE** | **P-Value** |
| **rs646776** | *SORT1; CELSR2; PSRC1* | Total Cholesterol | 4543 | -0.099 | 0.016 | $1.10 \times 10^{-9}$ |
| **rs646776** | *SORT1; CELSR2; PSRC1* | LDL Cholesterol | 4543 | -0.110 | 0.015 | $7.74 \times 10^{-14}$ |
| **rs646776** | *SORT1; CELSR2; PSRC1* | ApoB | 4546 | -2.695 | 0.328 | $2.66 \times 10^{-16}$ |
| **rs12129500** | *IL6R* | IL-6 | 4503 | -0.126 | 0.018 | $4.96 \times 10^{-12}$ |
| **rs11693654** | *ADCY3; NCOA1; CENPO* | BMI | 6387 | 0.200 | 0.036 | $3.57 \times 10^{-8}$ |
| **rs80026582** | *LPL* | Triglycerides | 4334 | -0.101 | 0.018 | $1.49 \times 10^{-8}$ |
| **rs80026582** | *LPL* | VLDL Cholesterol | 4334 | -0.100 | 0.018 | $1.57 \times 10^{-8}$ |
| **rs600038** | *ABO* | IL-6 | 4496 | -0.207 | 0.021 | $4.12 \times 10^{-22}$ |
| **rs174538** | *FADS1; FADS2; TMEM258* | Total Cholesterol | 4539 | -0.080 | 0.015 | $5.03 \times 10^{-8}$ |
| **rs2727784** | *APOA1; TAGLN* | ApoA1 | 4018 | 3.047 | 0.468 | $8.05 \times 10^{-11}$ |
| **rs10419998** | *GATAD2A; MAU2; TM6SF2* | ApoB | 4404 | -2.024 | 0.376 | $7.96 \times 10^{-8}$ |
| Abbreviations for the column headings from left to right: single nucleotide polymorphism, gene or gene cluster associated with SNP, associated trait, sample size for this effect, observed effect size (standard deviation), standard error of the effect size, p value for observed effect |  |  |  |  |  |  |

### Disentangling causal mechanisms using tissue-specific Mendelian randomization

After adjustment for the number of tests performed across all tissues and complex traits (p < 9.3×10^−4^ [p<0.05/54]), we identified 34 associations between tissue-specific gene expression and cardiovascular traits (Additional file 2: Tables S5-S15). In the validation analysis in ALSPAC, we observed consistent directions of effect for 30 of the associations. The potential value of this approach in terms of disentangling causal genes (i.e. scenarios 2 and 3) was exemplified at the BMI associated region on chromosome 2. Of the 3 cis- and potentially causal genes for this signal, only *ADCY3* provided strong evidence of being the putative causal gene in two types of adipose tissue (adipose subcutaneous (P = 6.8 × 10^−40^) and adipose visceral (P = 3.1 × 10^−48^)) (Figure 3a). This suggests that changes in *ADCY3* expression in adipose tissue could influence BMI levels. In contrast, there was a lack of evidence that changes in *NCOA1* expression in the analyzed tissue types influence BMI. We were unable to undertake MR of *CENPO* expression in this analysis as were unable to harmonise effect estimates between exposure and outcome. As an additional analysis, we repeated the MR on BMI using eQTL effect estimates derived from *ADCY3* expression in brain tissue (pituitary), although there was limited evidence of association (Beta (SE): 0.008 (0.006), P: 0.177).

**Figure 3.**
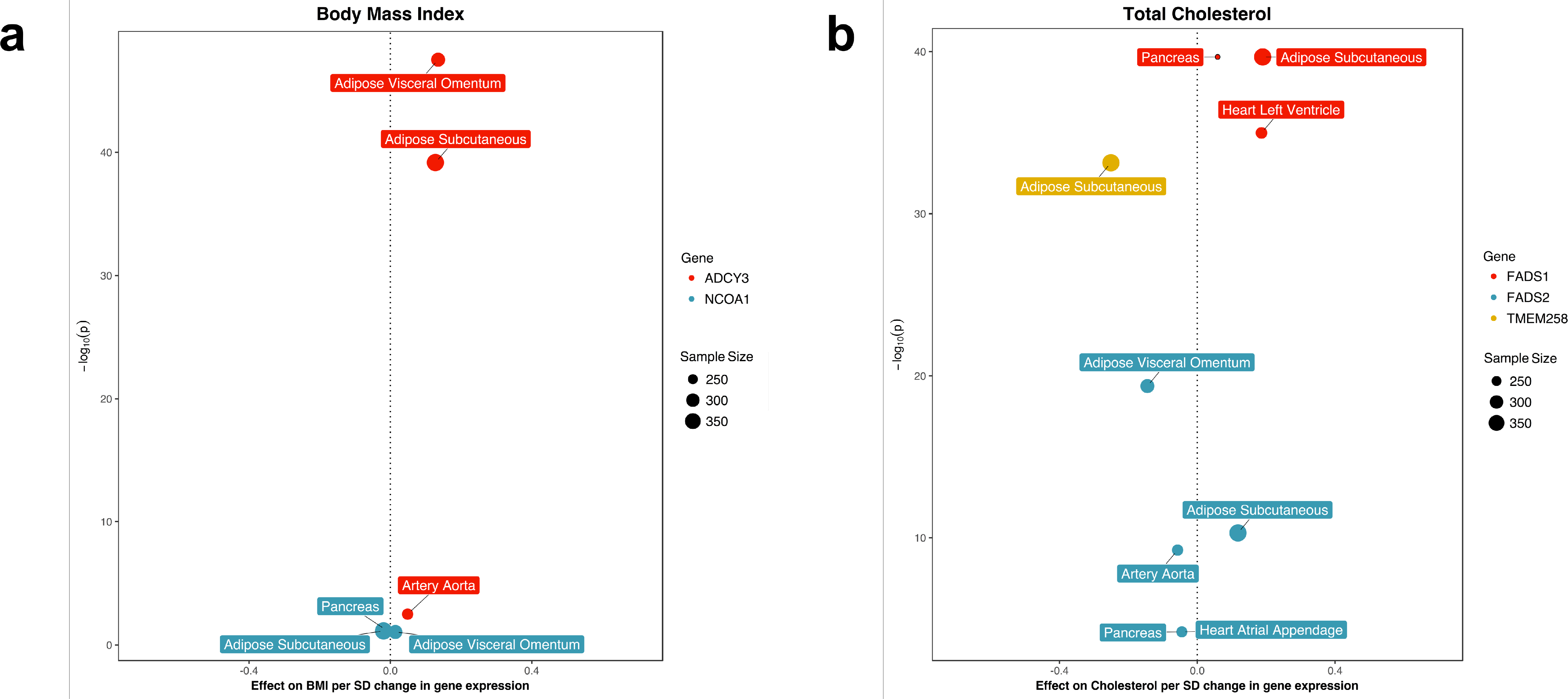
Volcano plots illustrating tissue-specific MR results. (a) Tissue-specific MR results for the observed effect on BMI. *ADCY3* gene expression provided strong evidence that it influenced BMI in comparison to the *NCOA1* gene. (b) Tissue-specific MR results for the observed effect on total cholesterol. All 3 genes provided strong evidence of association with total cholesterol at this region across various cardiovascular-specific tissue types.

Figure 3b illustrates results observed at the cholesterol associated region on chromosome 11. There was evidence that *FADS1* expression was associated with total cholesterol in 3 different tissues (adipose subcutaneous (P = 2.2 × 10^−40^), heart left ventricle (P = 1.0 × 10^−35^) and pancreas (P = 2.2 × 10^−40^)). Interestingly, the strength of evidence was comparable between subcutaneous adipose and pancreas tissues despite the differences in GTEx sample sizes (Pancreas: 220 & Adipose Subcutaenous: 385) (Additional file 1: Figure S2). *TMEM258* expression provided strong evidence of association in one tissue type (adipose subcutaneous (P = 7.2 × 10^−34^)), whereas association between *FADS2* expression and total cholesterol was observed in multiple tissue types (adipose subcutaneous (P = 5.1 × 10^−11^), adipose visceral (P = 4.2 × 10^−20^), artery aorta (P = 5.8 × 10^−10^), heart - atrial appendage (P = 6.3 × 10^−5^) and pancreas (P = 6.3 × 10^−5^)). The most parsimonious explanation may be that multiple genes at this locus influence cholesterol levels, however further analyses are required to robustly differentiate between scenarios 2 and 3 here (Figure 1).

At other loci evaluated (Additional File 1: Figure’s S3-S9), *LPL* showed evidence of association with triglycerides in a single tissue (adipose subcutaneous (P = 9.6 ×10^−168^)) implying that this effect may be more tissue-specific compared to those observed at other loci in this study (Additional file 1: Figure’s S8 & S9, Additional file 2: Tables S14 & S15). On chromosome 1, there was strong evidence that gene expression in liver influences total cholesterol (Additional file 1: Figure S6) and LDL (Additional file 1: Figure S7) (p < 3.22×10^−120^). However, this was observed for all three genes in the region (*SORT1*, *CELSR2* and *PSRC1*). In these analyses alone, we were unable to determine whether a particular gene is driving this observed effect, with the other proximal genes being coregulated, or whether there are multiple causal genes for these traits (i.e. scenario 2). However, evidence from the literature implicates *SORT1* as the most likely causal gene for this association signal [11,46]. Our MR results from ALSPAC provided evidence between *ABO* expression and IL-6 in 4 different tissues (Additional file 2: Table S12). Although, caution is required when interpreting this signal based on previous evidence across a diverse range of traits [47]. Finally, to test the direction of effect at each locus (i.e. are changes in gene expression causing changes in trait or vice versa), we ran a causal direction test [44]. In all scenarios, the test provided evidence that gene expression influences traits at these loci rather than the opposite direction of effect (Additional file 2: Tables S5-S15).

### Ascertaining whether DNA methylation resides on the causal pathway to disease

We identified evidence of colocalization (PPA ≥ 0.8) for 7 unique genes across 5 loci across various tissue types (Additional file 2: Tables S16-S20). Building upon results from the tissue-specific MR analysis, we found strong evidence that *ADCY3* is the functional gene for the BMI associated signal on chromosome 2 (maximum PPA of 0.99 between gene expression and BMI). We identified evidence of colocalization between BMI and *ADCY3* expression In both whole blood and subcutaneous adipose tissue. There was also evidence that distributions between DNA methylation at cg04553793 (at the promoter region of *ADCY3)* colocalized with BMI and *ADCY3* expression in whole blood (PPA = 0.88). However, the lead mQTL for this observed effect (rs13401333) was not correlated with the lead eQTL and GWAS hit (rs6745073, r^2^=0.02), which suggests that in-depth analysis with multiple tissue types is necessary to confirm whether DNA methylation influences disease suscepbility at this locus.

There was also evidence that changes in DNA methylation at a CpG site in the promoter region for *FADS1* (cg19610905) colocalized with total cholesterol variation. There was evidence of colocalization for all 3 traits using gene expression for *TMEM258* (PPA=0.85) (Figure4a), where the lead GWAS variant (rs174568) and mQTL were in perfect LD (rs1535, r^2^=1). This effect was only observed in whole blood. Evidence of colocalization between all three traits using *FADS1* expression narrowly missed the cut-off (PPA=0.77). Finally, we found limited evidence that changes in DNA methylation at this CpG site colocalized with *FADS2* expression, although as with the previously evaluated locus, this was not surprising given that cg19610905 is located downstream of *FADS2*. Gene expression of *TMEM258* in whole blood was negatively associated with DNA methylation at cg19610905. The directionality test suggested that DNA methylation influences TMEM258 expression at this locus rather than the opposite direction of effect (P<1 × 10^−16^).

**Figure 4.**
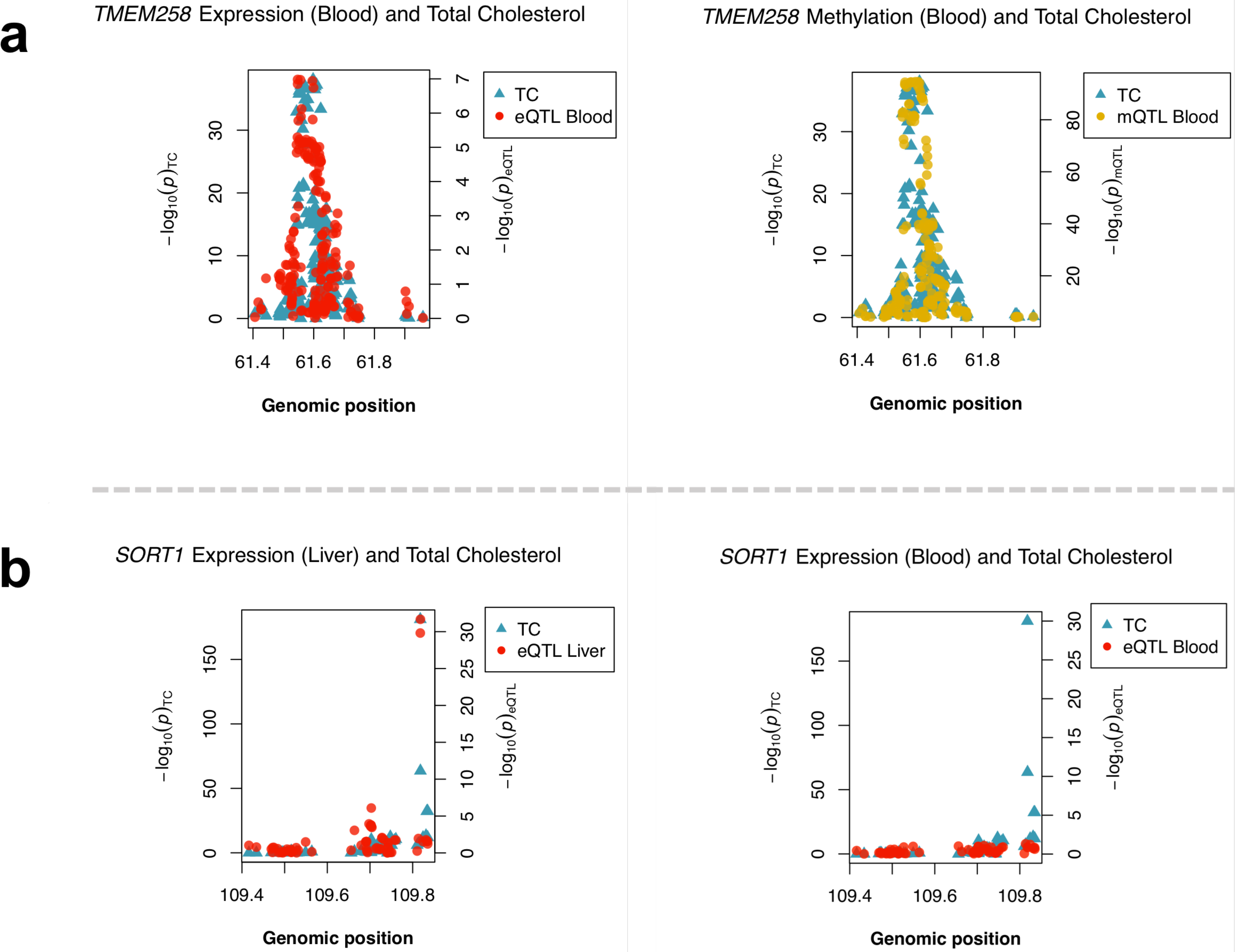
Multiple-trait colocalization analyses between cardiovascular traits and molecular phenotypes. (a) Evidence of colocalization between *TMEM258* expression and total cholesterol (left) as well as DNA methylation at cg19610905 and total cholesterol (right) using data derived from whole blood. Evidence of colocalization between *SORT1* expression using data derived from liver and total cholesterol (left). However, this evidence diminished when undertaking the same analysis for SORT1 expression data derived from whole blood (right).

We did not identify evidence in the colocalization analysis suggesting that DNA methylation plays a role in trait variation at the *SORT1* region. However, there was evidence of tissue specificity in liver tissue which supports evidence identified in our MR analysis. The first plot in Figure 4b illustrates how effects on *SORT1* gene expression and total cholesterol at this region colocalizes in liver tissue. In contrast, the neighbouring plot depicts the same analysis but in whole blood, whereby no evidence of colocalization was detected. Furthermore, we see the same tissue-specific colocalization for the effect on ApoB in the same region (Additional file 2: Table S16). The *CELSR2* gene showed similar evidence for tissue specificity in liver, whereas *PSRC1* expression colocalized with GWAS traits in both whole blood and liver.

## Discussion

In this study we have developed a framework to elucidate transcriptional mechanisms in disease which can help explain the functional relevance of GWAS findings. This is achieved by adapting the principles of MR to evaluating the putative effect of tissue-specific gene expression on complex traits, which can be complemented with moloc and harnessing large-scale summary statistics We demonstrate the value of this approach by evaluating 11 signals identified in a TWAS study undertaken in a cohort of young individuals from the ALSPAC cohort. Tissue-specific analyses helped infer whether individual or multiple genes were potentially responsible for observed signals at each locus. Moloc suggested that changes in gene expression and proximal DNA methylation may influence disease susceptibility at the *FADS1* locus.

The *ADCY3* locus has been reported to be associated with BMI in young individuals in previous studies [48,49]. Our MR analyses identified evidence that changes in *ADCY3* expression in adipose tissues may influence BMI, whereas weaker evidence was observed based on the expression of other proximal genes *(NCOA1)*. Specifically, we found that the magnitude of the effect for *ADCY3* expression was observed most strongly in adipose tissue, aligning with other research [50,51]. Furthermore, recent work has uncovered a variant in *ADCY3* associated with an increase in obesity levels [52]. In contrast, moloc showed a lack of evidence of colocalization for *NCOA1* expression. Moreover, although the *CENPO* gene was evaluated as part of our original association analysis, there were no eQTL for this gene for any of the tissues we analyzed. From this, we believe that *ADCY3* is likely the functional gene impacting BMI at this locus, although only with in-depth follow up analyses can this be determined with confidence. Our additional analysis indicated no tissue-specific effects using eQTL effect estimates derived from brain tissue, which suggests that the influence of *ADCY3* expression on BMI levels may be confined to adipose tissue. However, extended analyses using molecular data derived from brain tissue is necessary to confirm this, particularly given that previous work has linked gene expression in brain tissue with obesity-related traits [50,53].

We also identified evidence of colocalization for gene expression, DNA methylation and complex trait variation at the cholestrol associated region on chromosome 11. This was observed for *TMEM258* expression in whole blood, although *FADS1* narrowly missed the 0.8 cut-off (PPA = 0.77). This was based on DNA methylation levels at a CpG site located in the promoter region of *FADS1* (cg19610905). This effect was observed using data from whole blood (which is the only tissue we had accessible DNA methylation for in this study), which is potentially acting as a proxy for the true causal/relevant tissue type for this effect [18]. However, there was no indication that methylation played a role in the expression of *FADS2*. *TMEM258* has been proposed as a regulatory site for cholesterol in ‘*abdominal fat’* previously [54]. Interestingly, our MR analyses identified a single hit for this gene in adipose tissue, suggesting that *TMEM258* expression is highly tissue-specific. *FADS1* has previously been associated with cholesterol levels in young individuals [55]. Additionally, genetic variation at this region is associated with DNA methylation levels at cg19610905 based on cord blood in ARIES, which suggests that these methylation changes may influence the expression of *FADS1/TMEM258* from a very early age. Overall at this region, our results suggest that scenario 2 is a likely explanation for the association signal, where it is biologically plausible that multiple causal genes influence complex trait variation. Specifically, our analyses suggest that *TMEM258* and *FADS1* are potential causal genes, however, further work is needed to elucidate whether *FADS2* is directly influencing cardiovascular traits or is simply co-regulated with the nearby functional loci.

The *LPL* locus was not subject to co-regulation/uncertainty over the likely causal gene and is therefore likely attributed to scenario 1. *LPL* has been previously reported to influence lipid and triglyceride levels [56–58] and there is also evidence from gene knockout experiments [59]. The tissue-specificity of *LPL* has also previously been explored, although not by recent studies [60]. 2SMR analyses provided robust evidence of highly specific gene expression in adipose tissue, corroborating previous research [60,61].

For other regions evaluated in our study, there was evidence that multiple genes may potentially influence traits. The *SORT1* locus has been previously studied in detail with regards to its effect on cholesterol levels [46,62]. Our MR analyses provided additional evidence of an effect using expression derived from liver tissue for *SORT1*, *CELSR2* and *PSRC1*, as well as in pancreas tissue for *SORT1* and *CELSR2* only. Our subsequent moloc analysis identified evidence of colocalization for *SORT1* and *CELSR2* expression with cholesterol only in liver tissue, suggesting that *PSRC1* could be less tissue-specific than the other 2 genes in this region. Previous research supports these observations with regards to the effects of *SORT1* and *CELSR2* in liver [11,63], as well as the lack of tissue-specificity for the *PSRC1* locus [64]. There was limited evidence that DNA methylation was affecting gene expression at this region, although future work with methylation data derived from liver tissue is warranted.

This study has demonstrated the value of our systematic framework in terms of distinguishing between scenarios 1, 2, 3 and 4. However, an important limiting factor, as with any study applying single-instrument MR, is the inability to separate mediation from horizontal pleiotropy (i.e. scenario 5). Given that *trans*-eQTLs likely regulate genes through a non-allele-specific mechanism [65], we selected only eQTLs that were influencing proximal genes. As more eQTL are uncovered across the genome by future studies, across a wide range of tissue and cell types, our framework should become increasingly powerful to evaluate all 5 outlined scenarios.

In terms of limitations in this study, we recognise the varying sample sizes between tissues in GTEx will determine the relative power to detect eQTL (Additional file 1: Figure S2). Increased sample sizes in GTEx [66] and similar endeavours will help address this limitation. Furthermore, the DNA methylation data we incorporated within our framework from the accessible resource for ARIES [45] project was only obtained in whole blood. However, in general, investigating the potential mediatory role of DNA methylation in whole blood is a limitation, as this assumes that whole blood is acting as a proxy for another, more relevant tissue type [67]. Furthermore, recent work has suggested that promoter DNA methylation may not be sufficient on its own to influence transcriptional changes [68]. Future work will need to incorporate DNA methylation data from various tissues as and when these data become available so we can better understand the role of this epigenetic process on transcriptional activity. For this purpose, a resource concerning tissue-specific DNA methylation would be extremely valuable.

Another constraint of relatively modest sample sizes in GTEx is that we did not detect evidence of co-localization at some loci despite investigating the functionally relevant gene. For example, we can be reasonably certain that circulating ApoA1 levels are influenced by the expression of *APOA1*. The complexity of gene regulation is often underestimated due to factors such as feedback loops, hidden confounders in expression data and regulatory activity not always being detected in relevant tissues [69]. However, we are beginning to better understand regulation across tissues [64], which should provide us with further opportunities to detect cross-tissue regulatory activity and develop our biological understanding of disease.

## Conclusions

We have identified a number of tissue-specific effects at several regions throughout the genome. Our results suggest that DNA methylation may also influence complex traits through gene expression pathways for observed effects on BMI and cholesterol. In-depth evaluations of the loci identified in our study should help fully understand the causal pathway to disease for these effects. Furthermore, as these genetic loci influence cardiovascular traits early in the life course, these endeavours should allow a long window of intervention for disease susceptibility. Finally, the framework outlined in this study should prove particularly valuable for future studies as increasingly large datasets concerning tissue-specific gene expression become available.

## Abbreviations

GWAS: Genome-wide association study
LDL: Low-density lipoprotein
SNP: Single nucleotide polymorphism
mRNA: Messenger ribonucleic acid
eQTL: Expression quantitative trait loci
Mb: Megabase
TSS: Transcription start site
TWAS: Transcription-wide association study
MR: Mendelian Randomization
GTEx: Genotype tissue expression project
LD: Linkage Disequilibrium
Moloc: Multiple-trait colocalization
mQTL: Methylation quantitative trait loci
ALSPAC: Avon Longitudinal Study of Parents and Children
MAF: Minor allele frequency
BMI: Body mass index
VLDL: Very low-density lipoprotein
ApoA1: Apolipoprotein A1
ApoB: Apolipoprotein B
CRP: C-Reactive protein
IL-6: Interleukin 6
ARIES: Accessible resource for integrated epigenomic studies
PPA: Posterior probability of association

## Declarations

### Ethics approval and consent to participate

All procedures were ethically approved by the ALSPAC ethics and Law Committee and the Local Research Ethics Committees. Written informed consent was obtained from all participants. We were granted access to ALSPAC data under project B2965 “Evaluating the causal effect of gene expression on cardiovascular function” (04/10/2017).

### Consent for publication

This project has been approved for publication by the ALSPAC executive committee.

### Availability of data and material

Access to ALSPAC and ARIES data is available to all bona fide researchers submitting a research proposal at www.bristol.ac.uk/alspac. GTEx (https://www.gtexportal.org/home/) and large-scale GWAS data (refer to Additional file 2:Table S4) is publicly available data which does not require a proposal for access.

### Authors contributions

TGR led the design of the project. TGR and TRG supervised the project. KT undertook statistical and bioinformatics analysis. KT and TGR drafted the manuscript. Comments were provided by TRG, GDS and CLR. All authors approved the final version of the manuscript.

## Acknowledgements

We are grateful to everyone involved in the Avon Longitudinal Study of Parents and Children (ALSPAC). This includes the families who kindly participated, the midwives for recruiting them and everyone behind the scenes ensuring the smooth running of the study. The UK Medical Research Council, the Wellcome Trust (grant 102215/2/13/2) and the University of Bristol provide core support for ALSPAC. The author greatly appreciates the help received from Christian Benner (lead developer of FINEMAP) as well as Claudia Giambartolomei (lead developer of moloc) for her help with the multiple trait colocalization methods. This work was supported by the British Heart Foundation [SSCM SJ1429] and UK Medical Research Council (MC_UU_12013/8). TGR is a UKRI Innovation Research Fellow (MR/S003886/1).

## Additional files

### Additional file 1 - Supplementary figures

**Figure S1**. MR effect estimates are based on the Wald ratio test, where 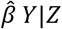 is the coefficient of the genetic variant in the regression of the exposure (e.g. gene expression) and 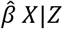 is the coefficient of the genetic variant in the regression of the outcome (e.g. cardiovascular trait). **Figure S2**. Scatter plot illustrating how eGene discovery increases as sample size increases (R^2^ = 0.84). Figure adapted from the Genotype Tissue Expression Project (Aguet et al 2017). **Figure S3**. Volcano plot from our tissue-specific Mendelian randomization analysis for the Apolipoprotein A1 associated region (rs2727784). Outcome data from Kettunen et al (2016). **Figure S4**. Volcano plot from our tissue-specific Mendelian randomization analysis for the Apolipoprotein B associated region (rs646776). Outcome data from Kettunen et al (2016). **Figure S5**. Volcano plot from our tissue-specific Mendelian randomization analysis for the Apolipoprotein B associated region (rs10419998). Outcome data from Kettunen et al (2016). **Figure S6**. Volcano plot from our tissue-specific Mendelian randomization analysis for the cholesterol associated region (rs646776). Outcome data from Willer CJ et al (2016). **Figure S7**. Volcano plot from our tissue-specific Mendelian randomization analysis for the low density lipoprotein associated region (rs646776). Outcome data from Willer CJ et al (2016). **Figure S8**. Volcano plot from our tissue-specific Mendelian randomization analysis for the triglyceride associated region (rs80026582). Outcome data from Willer CJ et al (2016). **Figure S9**. Volcano plot from our tissue-specific Mendelian randomization analysis for the very low density lipoprotein associated region (rs80026582). Outcome data from Kettunen et al (2016).

### Additional file 2 - Supplementary tables

**Table S1**. Tissues used for tissue-specific Mendelian randomization. **Table S2**. Results of fine mapping analysis. **Table S3**. Tissues used for moloc analysis. **Table S4**. Details on the GWAS datasets used. **Table S5**. Tissue-specific Mendelian Randomization results for the Apoliporotein A1 associated region on chromosome 11 (rs2727784). **Table S6**. Tissue-specific Mendelian randomization results for the Apolipoprotein B associated region on chromosome 1 (rs646776). **Table S7**. Tissue-specific Mendelian randomization results for the Apolipoprotein B associated region on chromosome 19 (rs10419998). **Table S8**. Tissue-specific Mendelian randomization results for the body mass index associated region chromosome 2 (rs11693654). **Table S9**. Tissue-specific Mendelian randomization results for the cholesterol associated region on chromosome 1 (rs646776). **Table S10**. Tissue-specific Mendelian randomization results for the cholesterol associated region on chromosome 11 (rs174538). **Table S11**. Tissue-specific Mendelian randomization results for the interleukin-6 associated region on chromosome 1 (rs12129500). **Table S12**. Tissue-specific Mendelian randomization results for the interleukin-6 associated region on chromosome 9 (rs600038).**Table S13**. Tissue-specific Mendelian randomization results for the low density lipoprotein associated region on chromosome 1 (rs646776). **Table S14**. Tissue-specific Mendelian randomization results for the triglyceride associated region on chromosome 8 (rs80026582). **Table S15**. Tissue-specific Mendelian randomization results for the very low density lipoprotein associated region on chromosome 8 (rs80026582). **Table S16**. Moloc results for the apolipoprotein B associated region on chromosome 1. **Table S17**. Moloc results for the cholesterol associated region on chromosome 1. **Table S18**. Moloc results for the body mass index associated region on chromosome 2. **Table S19**. Moloc results for the cholesterol associated region on chromosome 11. **Table S20**. Moloc results for the low density lipoprotein associated region on chromosome 1.

## References

1. World Health Organization. Cardiovascular Disease: Global Atlas on Cardiovascular Disease Prevention and Control. Geneva, Switzerland; 2012.

2. Altshuler D, Daly MJ, Lander E. Genetic Mapping in Human Disease. Science (80-). 2009;322:881–8.

3. Smith JG, Newton-Cheh C. Genome-wide association studies of late-onset cardiovascular disease. J. Mol. Cell. Cardiol. 2015. p. 131–41.

4. Holmes M V., Asselbergs FW, Palmer TM, Drenos F, Lanktree MB, Nelson CP, et al. Mendelian randomization of blood lipids for coronary heart disease. Eur Heart J. 2015;36:539–50.

5. Mihaylova B, Emberson J, Blackwell L, Keech A, Simes J, Barnes EH, et al. The effects of lowering LDL cholesterol with statin therapy in people at low risk of vascular disease: meta-analysis of individual data from 27 randomised trials. Lancet [Internet]. 2012;380:581–90. Available from: http://www.pubmedcentral.nih.gov/articlerender.fcgi?artid=3437972&tool=pmcentrez&rendertype=abstract%0A http://www.sciencedirect.com/science/article/pii/S0140673612603675

6. Hindorff LA, Sethupathy P, Junkins HA, Ramos EM, Mehta JP, Collins FS, et al. Potential etiologic and functional implications of genome-wide association loci for human diseases and traits. Proc Natl Acad Sci [Internet]. 2009;106:9362–7. Available from: http://www.pnas.org/cgi/doi/10.1073/pnas.0903103106

7. Edwards SL, Beesley J, French JD, Dunning M. Beyond GWASs: Illuminating the dark road from association to function. Am. J. Hum. Genet. 2013. p. 779–97.

8. Joehanes R, Zhang X, Huan T, Yao C, Ying S, Nguyen QT, et al. Integrated genome-wide analysis of expression quantitative trait loci aids interpretation of genomic association studies. Genome Biol [Internet]. 2017;18:16. Available from: http://genomebiology.biomedcentral.com/articles/10.1186/s13059-016-1142-6

9. Gusev A, Ko A, Shi H, Bhatia G, Chung W, Penninx BWJH, et al. Integrative approaches for large-scale transcriptome-wide association studies. Nat Genet [Internet]. Nature Publishing Group; 2016;48:245–52. Available from: http://www.nature.com/doifinder/10.1038/ng.3506

10. Nica AC, Dermitzakis ET. Expression quantitative trait loci: present and future. Philos Trans R Soc Lond B Biol Sci [Internet]. 2013;368:2012–0362. Available from: http://www.ncbi.nlm.nih.gov/pubmed/23650636%5Cnhttp://www.pubmedcentral.nih.gov/articlerender.fcgi?artid=PMC3682727%5Cnhttp://rstb.royalsocietypublishing.org/content/368/1620/20120362

11. Wainberg M, Sinnott-Armstrong N, Knowles D, Golan D, Ermel R, Ruusalepp A, et al. Vulnerabilities of transcriptome-wide association studies. bioRxiv [Internet]. 2017; Available from: http://biorxiv.org/content/early/2017/10/20/206961.abstract

12. Davey Smith G, Hemani G. Mendelian randomization: genetic anchors for causal inference in epidemiological studies. Hum Mol Genet [Internet]. 2014;23:R89–98. Available from: http://www.ncbi.nlm.nih.gov/pubmed/25064373%5Cnhttp://www.pubmedcentral.nih.gov/articlerender.fcgi?artid=PMC4170722

13. Davey Smith G, Ebrahim S. “Mendelian randomization”:Can genetic epidemiology contribute to understanding environmental determinants of disease? Int J Epidemiol. 2003;32:1–22.

14. Lawlor DA. Commentary: Two-sample Mendelian randomization: opportunities and challenges. Int J Epidemiol [Internet]. 2016;45:908–15. Available from: https://academic.oup.com/ije/article-lookup/doi/10.1093/ije/dyw127

15. Aguet F, Ardlie KG, Cummings BB, Gelfand ET, Getz G, Hadley K, et al. Genetic effects on gene expression across human tissues. Nature [Internet]. 2017;550:204–13. Available from: http://www.nature.com/doifinder/10.1038/nature24277

16. Giambartolomei C, Zhenli Liu J, Zhang W, Hauberg M, Shi H, Boocock J, et al. A Bayesian Framework for Multiple Trait Colocalization from Summary Association Statistics. bioRxiv [Internet]. 2017; Available from: http://biorxiv.org/content/early/2017/06/26/155481.abstract

17. Wahl S, Drong A, Lehne B, Loh M, Scott WR, Kunze S, et al. Epigenome-wide association study of body mass index, and the adverse outcomes of adiposity. Nature [Internet]. 2016;541:81–6. Available from: http://www.nature.com/doifinder/10.1038/nature20784

18. Qi T, Wu Y, Zeng J, Zhang F, Xue A, Jiang L, et al. Identifying gene targets for brain-related traits using transcriptomic and methylomic data from blood. Nat Commun. 2018;1–26.

19. Bonder MJ, Luijk R, Zhernakova D V., Moed M, Deelen P, Vermaat M, et al. Disease variants alter transcription factor levels and methylation of their binding sites. Nat Genet. 2017;49:131–8.

20. Acharya CR, Owzar K, Allen AS. Mapping eQTL by leveraging multiple tissues and DNA methylation. BMC Bioinformatics. 2017; 18.

21. Hannon E, Dempster E, Viana J, Burrage J, Smith AR, Macdonald R, et al. An integrated genetic-epigenetic analysis of schizophrenia: Evidence for co-localization of genetic associations and differential DNA methylation. Genome Biol. 2016;17.

22. Hodgkin J. Seven types of pleiotropy. Int. J. Dev. Biol. 1998. p. 501–5.

23. Hong YM. Atherosclerotic cardiovascular disease beginning in childhood. Korean Circ. J. 2010. p. 1–9.

24. Golding J, Pembrey M, Jones R. ALSPAC--the Avon Longitudinal Study of Parents and Children. I. Study methodology. Paediatr Perinat Epidemiol. 2001;15:74–87.

25. Fraser A, Macdonald-wallis C, Tilling K, Boyd A, Golding J, Davey smith G, et al. Cohort profile: The avon longitudinal study of parents and children: ALSPAC mothers cohort. Int J Epidemiol. 2013;42:97–110.

26. Boyd A, Golding J, Macleod J, Lawlor DA, Fraser A, Henderson J, et al. Cohort profile: The ‘Children of the 90s’-The index offspring of the avon longitudinal study of parents and children. Int J Epidemiol. 2013;42:111–27.

27. University of Bristol. Accessing the resource [Internet]. [cited 2018 Jan 29]. Available from: http://www.bristol.ac.uk/alspac/researchers/access/

28. 1000 Genomes Project Consortium, Auton A, Brooks LD, Durbin RM, Garrison EP, Kang HM, et al. A global reference for human genetic variation. Nature [Internet]. 2015;526:68–74. Available from: http://www.ncbi.nlm.nih.gov/pubmed/26432245%0Ahttp://www.pubmedcentral.nih.gov/articlerender.fcgi?artid=PMC4750478

29. McCarthy S, Das S, Kretzschmar W, Delaneau O, Wood AR, Teumer A, et al. A reference panel of 64,976 haplotypes for genotype imputation. Nat Genet. 2016;48:1279–83.

30. Warnick GR, Knopp RH, Fitzpatrick V, Branson L. Estimating low-density lipoprotein cholesterol by the Friedewald equation is adequate for classifying patients on the basis of nationally recommended cutpoints. Clin Chem. 1990;36:15–9.

31. Falaschetti E, Hingorani AD, Jones A, Charakida M, Finer N, Whincup P, et al. Adiposity and cardiovascular risk factors in a large contemporary population of prepubertal children. Eur Heart J. 2010;31:3063–72.

32. Westra HJ, Peters MJ, Esko T, Yaghootkar H, Schurmann C, Kettunen J, et al. Systematic identification of trans eQTLs as putative drivers of known disease associations. Nat Genet. 2013;45:1238–43.

33. Conesa A, Madrigal P, Tarazona S, Gomez-Cabrero D, Cervera A, McPherson A, et al. A survey of best practices for RNA-seq data analysis. Genome Biol. 2016.

34. Chang CC, Chow CC, Tellier LCAM, Vattikuti S, Purcell SM, Lee JJ. Second-generation PLINK: Rising to the challenge of larger and richer datasets. Gigascience. 2015;4.

35. Purcell S, Chang C. PLINK 1.9 [Internet]. 2015 [cited 2018 Jan 9]. Available from: href="www.cog-genomics.org/plink/1.9/

36. Turner SD. qqman: an R package for visualizing GWAS results using Q-Q and manhattan plots [Internet]. bioRxiv. 2014. Available from: http://biorxiv.org/lookup/doi/10.1101/005165

37. Benner C, Spencer CCA, Havulinna AS, Salomaa V, Ripatti S, Pirinen M. FINEMAP: Efficient variable selection using summary data from genome-wide association studies. Bioinformatics. 2016;32:1493–501.

38. Burgess S, Small DS, Thompson SG. A review of instrumental variable estimators for Mendelian randomization. Stat. Methods Med. Res. 2017.

39. Kettunen J, Demirkan A, Würtz P, Draisma HHM, Haller T, Rawal R, et al. Genome-wide study for circulating metabolites identifies 62 loci and reveals novel systemic effects of LPA. Nat Commun. 2016;7.

40. Willer CJ, Schmidt EM, Sengupta S, Peloso GM, Gustafsson S, Kanoni S, et al. Discovery and refinement of loci associated with lipid levels. Nat Genet. 2013;45:1274–85.

41. Bycroft C, Freeman C, Petkova D, Band G, Elliott LT, Sharp K, et al. Genome-wide genetic data on ~500,000 UK Biobank participants. bioRxiv [Internet]. 2017; Available from: http://biorxiv.org/content/early/2017/07/20/166298.abstract

42. Hemani G, Zheng J, Wade KH, Laurin C, Elsworth B, Burgess S, et al. MR-Base: a platform for systematic causal inference across the phenome using billions of genetic associations. bioRxiv [Internet]. 2016;078972. Available from: https://www.biorxiv.org/content/early/2016/12/16/078972

43. Wickham H. ggplot2 Elegant Graphics for Data Analysis [Internet]. Media. 2009. Available from: http://had.co.nz/ggplot2/book

44. Hemani G, Tilling K, Davey Smith G. Orienting the causal relationship between imprecisely measured traits using GWAS summary data. PLoS Genet. 2017;13.

45. Relton CL, Gaunt T, McArdle W, Ho K, Duggirala A, Shihab H, et al. Data resource profile: Accessible resource for integrated epigenomic studies (ARIES). Int J Epidemiol. 2015;44:1181–90.

46. Musunuru K, Strong A, Frank-Kamenetsky M, Lee NE, Ahfeldt T, Sachs K V., et al. From noncoding variant to phenotype via SORT1 at the 1p13 cholesterol locus. Nature. 2010;466:714–9.

47. Pickrell JK, Berisa T, Liu JZ, Ségurel L, Tung JY, Hinds DA. Detection and interpretation of shared genetic influences on 42 human traits. Nat Genet. 2016;48:709–17.

48. Stergiakouli E, Gaillard R, Tavaré JM, Balthasar N, Loos RJ, Taal HR, et al. Genome-wide association study of height-adjusted BMI in childhood identifies functional variant in ADCY3. Obesity. 2014;22:2252–9.

49. Namjou B, Keddache M, Marsolo K, Wagner M, Lingren T, Cobb B, et al. EMR-linked GWAS study: Investigation of variation landscape of loci for body mass index in children. Front Genet. 2013;4.

50. Hao R-H, Yang T-L, Rong Y, Yao S, Dong S-S, Chen H, et al. Gene expression profiles indicate tissue-specific obesity regulation changes and strong obesity relevant tissues. Int J Obes [Internet]. 2018;1–7. Available from: http://www.nature.com/doifinder/10.1038/ijo.2017.283

51. Vink RG, Roumans NJ, Fazelzadeh P, Tareen SH, Boekschoten M V, van Baak MA, et al. Adipose tissue gene expression is differentially regulated with different rates of weight loss in overweight and obese humans. Int J Obes [Internet]. 2017;41:309–16. Available from: https://www.ncbi.nlm.nih.gov/pubmed/27840413

52. Grarup N, Moltke I, Andersen MK, Dalby M, Vitting-Seerup K, Kern T, et al. Loss-of-function variants in ADCY3 increase risk of obesity and type 2 diabetes. Nat Genet. 2018;

53. Samad F, Pandey M, Loskutoff DJ. Regulation of tissue factor gene expression in obesity. Blood. 2001;98:3353–8.

54. Franzén O, Ermel R, Cohain A, Akers NK, Di Narzo A, Talukdar HA, et al. Cardiometabolic risk loci share downstream cis- and trans-gene regulation across tissues and diseases. Science (80-). 2016;353:827–30.

55. Dumont J, Huybrechts I, Spinneker A, Gottrand F, Grammatikaki E, Bevilacqua N, et al. FADS1 Genetic Variability Interacts with Dietary -Linolenic Acid Intake to Affect Serum Non-HDL-Cholesterol Concentrations in European Adolescents. J Nutr [Internet]. 2011;141:1247–53. Available from: http://jn.nutrition.org/cgi/doi/10.3945/jn.111.140392

56. Johansen CT, Kathiresan S, Hegele RA. Genetic determinants of plasma triglycerides. J Lipid Res [Internet]. 2011;52:189–206. Available from: http://www.jlr.org/lookup/doi/10.1194/jlr.R009720

57. Humphries SE, Nicaud V, Margalef J, Tiret L, Talmud PJ. Lipoprotein lipase gene variation is associated with a paternal history of premature coronary artery disease and fasting and postprandial plasma triglycerides: the European Atherosclerosis Research Study (EARS). Arterioscler Thromb Vasc Biol. 1998;18:526–34.

58. Mead JR, Irvine S a, Ramji DP. Lipoprotein lipase: structure, function, regulation, and role in disease. J Mol Med (Berl) [Internet]. 2002;80:753–69. Available from: http://www.ncbi.nlm.nih.gov/pubmed/12483461

59. Chen Y, Zhu J, Lum PY, Yang X, Pinto S, MacNeil DJ, et al. Variations in DNA elucidate molecular networks that cause disease. Nature. 2008;452:429–35.

60. Ranganathan G, Ong JM, Yukht A, Saghizadeh M, Simsolo RB, Pauer A, et al. Tissue-specific expression of human lipoprotein lipase: Effect of the 3???-untranslated region on translation. J Biol Chem. 1995;270:7149–55.

61. Wang H, Eckel RH. Lipoprotein lipase: from gene to obesity. Am J Physiol Endocrinol Metab [Internet]. 2009;297:E271–88. Available from: http://ajpendo.physiology.org/content/297/2ZE271

62. Arvind P, Nair J, Jambunathan S, Kakkar V V., Shanker J. CELSR2-PSRC1-SORT1 gene expression and association with coronary artery disease and plasma lipid levels in an Asian Indian cohort. J Cardiol. 2014;64:339–46.

63. Schadt EE, Molony C, Chudin E, Hao K, Yang X, Lum PY, et al. Mapping the genetic architecture of gene expression in human liver. PLoS Biol. 2008;6:1020–32.

64. Ongen H, Brown AA, Delaneau O, Panousis NI, Nica AC, Dermitzakis ET. Estimating the causal tissues for complex traits and diseases. Nat Genet. 2017;49:1676–83.

65. Albert FW, Kruglyak L. The role of regulatory variation in complex traits and disease. Nat. Rev. Genet. 2015. p. 197–212.

66. Stranger BE, Brigham LE, Hasz R, Hunter M, Johns C, Johnson M, et al. Enhancing GTEx by bridging the gaps between genotype, gene expression, and disease. Nat. Genet. 2017. p. 1664–70.

67. Teschendorff AE, Relton CL. Statistical and integrative system-level analysis of DNA methylation data. Nat Rev Genet [Internet]. 2017; Available from: http://www.nature.com/doifinder/10.1038/nrg.2017.86

68. Ford EE, Grimmer MR, Stolzenburg S, Bogdanovic O, de Mendoza A, Farnham PJ, et al. Frequent lack of repressive capacity of promoter DNA methylation identified through genome-wide epigenomic manipulation. bioRxiv [Internet]. 2017; Available from: http://biorxiv.org/content/early/2017/09/20/170506.abstract

69. Torres JM, Barbeira AN, Bonazzola R, Morris AP, Shah KP, Wheeler HE, et al. Integrative cross tissue analysis of gene expression identifies novel type 2 diabetes genes. bioRxiv [Internet]. 2017;108–134. Available from: http://biorxiv.org/content/early/2017/02/27/108134

